# AOP Net: An AI-Enhanced Software Platform for the Visualization and Analysis of Complex Toxicological Pathways

**DOI:** 10.1101/2025.08.25.672239

**Authors:** Abhik Seal

**Author notes:** Use footnote for providing further information about author (webpage, alternative address)—*not* for acknowledging funding agencies.

## Abstract

The Adverse Outcome Pathway (AOP) framework is a cornerstone of 21st-century toxicology, providing a structured method for organizing mechanistic knowledge to support risk assessment. However, the inherent complexity of biological systems, characterized by interconnected signaling networks, reveals the limitations of simple, linear AOP representations. This has created a pressing need for advanced computational tools that can visualize and analyze these intricate AOP networks. This study introduces the AOP Network Viewer, a novel software platform designed to address the critical challenges of scalability, interactivity, and deep analysis in toxicological research. The system is built on a scalable client-server architecture with a graph database backend, supporting sophisticated interactive features such as multi-pathway selection, dynamic path-finding, and advanced filtering. The principal innovation of the AOP Network Viewer is its integrated AI Analysis Panel, a Large Language Model (LLM)-powered engine that provides a deep, context-aware analysis of selected network components. This AI system utilizes a unique three-layer prompt architecture to deliver domain-specific insights across seven specialized analytical modes. Case studies demonstrate the platform’s utility in developing alternative testing strategies for endocrine disruptors, interpreting high-throughput screening data, and generating hypotheses for toxicity assessment of mixtures. The AOP Network Viewer is engineered to translate complex user queries into focused, interpretable biological networks, transforming AOP network visualization from a passive viewing experience into an interactive and intelligent analytical process, thereby accelerating mechanistic discovery and enhancing evidence-based regulatory decision-making. The app is source code is available at https://github.com/abhik1368/aop-visualizer-clean.

## 1 Introduction

The field of toxicology has been undergoing a profound paradigm shift, moving away from a primary reliance on traditional, apical endpoint-based animal testing towards a more mechanistic, human-relevant approach to safety and risk assessment.1 This transformation was envisioned by the U.S. The National Research Council’s 2007 report, “Toxicity Testing in the 21st Century,” is driven by the ethical, economic, and scientific limitations of conventional methods in addressing the vast number of chemicals requiring evaluation. At the heart of this evolution is the Adverse Outcome Pathway (AOP) framework U.S. Environmental Protection Agency [2017], Knapen et al. [2018] first conceptualized by Ankley et al. in 2010.

An adverse outcome pathway (AOP) is essentially a roadmap that shows how a chemical or environmental stressor can lead to harmful effects in an organism. It is built from a series of measurable steps called Key Events (KEs). This chain reaction starts with a Molecular Initiating Event (MIE), which is the first point of contact between the stressor and a molecule in the body. This trigger leads to a cascade of intermediate KEs that progress from the cellular level to the organ or individual level. The sequence ends with the final Adverse Outcome (AO), which could be a disease in an individual or a population decline in an ecosystem. The links connecting each of these steps are called Key Event Relationships (KERs), which describe the cause-and-effect relationship between one event and the next.

To make them useful and efficient, AOPs have been developed based on a few key principles. They are designed to be general and not specific to any single chemical, allowing one pathway to be applied to many different substances. The components, such as the individual Key Events, are modular, meaning they can be reused in different AOPs, similar to building blocks. While each AOP is developed as a single, linear path, their real value comes when they are combined into complex networks that better reflect actual biology. Finally, AOPs are considered “living documents.” They are not set in stone and can be updated and refined as new scientific evidence becomes available, ensuring that they remain accurate and relevant for assessing risks.

The AOP framework provides a structured, transparent, and collaborative approach to organizing existing biological and toxicological knowledge into a simple causal sequence [1]. This structure enables the integration of data from diverse sources, including in vitro assays and computational models, providing a clear mechanistic line of sight from chemical exposure to adverse health effects.

### 1.1 From Linear Pathways to Complex Networks

While the linear representation of an individual AOP is conceptually elegant and pragmatically useful for development and documentation, it is a deliberate oversimplification of the biological reality. Biological systems do not operate as isolated linear chains of events. Instead, they are characterized by dense webs of interconnected pathways featuring crosstalk, feedback loops, and redundancy. A single chemical stressor can trigger multiple MIEs, activating numerous pathways that may converge on a common KE or diverge to produce a spectrum of AOs. This biological reality has necessitated a critical evolution in the AOP framework, specifically the transition from individual linear pathways to complex AOP networks. An AOP network is defined as an assembly of two or more AOPs that share at least one common component, typically a KE. This network perspective allows for a more realistic and powerful representation of toxicology, capturing phenomena such as :

- Convergent Networks: Multiple distinct upstream pathways leading to a common AO, which is critical for understanding how different chemical classes can produce similar toxicities.
- Divergent Networks: A single MIE or early KE leading to multiple distinct AOs, explaining the pleiotropic effects of certain chemicals.
- Mixed Networks: The most biologically realistic representation, combining convergent and divergent elements, often forming “bow-tie” motifs, where numerous inputs are integrated at critical hub KEs before branching out to various outcomes.

The modular design of the AOP-Wiki, the central knowledge base for AOPs curated under the guidance of the Organization for Economic Co-operation and Development (OECD), inherently supports this network concept by allowing KEs to be shared across multiple AOPs. As the AOP-Wiki grows, these networks emerge organically, creating a rich but complex tapestry of mechanistic insights.

Figure 1. The Adverse Outcome Pathway (AOP) network illustrates the complex pathogenesis of liver fibrosis, highlighting how disparate Molecular Initiating Events (MIEs) can converge through interconnected pathways to a common adverse outcome. The network is initiated by at least three distinct MIEs: activation of the Aryl Hydrocarbon Receptor (AhR), covalent protein alkylation, and endocytotic lysosomal uptake. These triggers induce a cascade of early Key Events (KEs), with protein alkylation and endocytotic uptake directly converging on the induction of cell injury and death. Concurrently, endocytic uptake can lead to lysosomal disruption, which, along with AhR activation, contributes to mitochondrial dysfunction, further exacerbating cellular damage. This widespread cell injury serves as a critical node, instigating a proinflammatory response characterized by the release of mediators and leukocyte recruitment. Ultimately, these cytotoxic and inflammatory cascades converge on the pivotal KE: the activation of hepatic stellate cells. Once activated, these cells drive the subsequent KE of increased extracellular matrix deposition, which is the direct precursor to the final adverse outcome of liver fibrosis Granzow et al. [2014].

**Figure 1:**
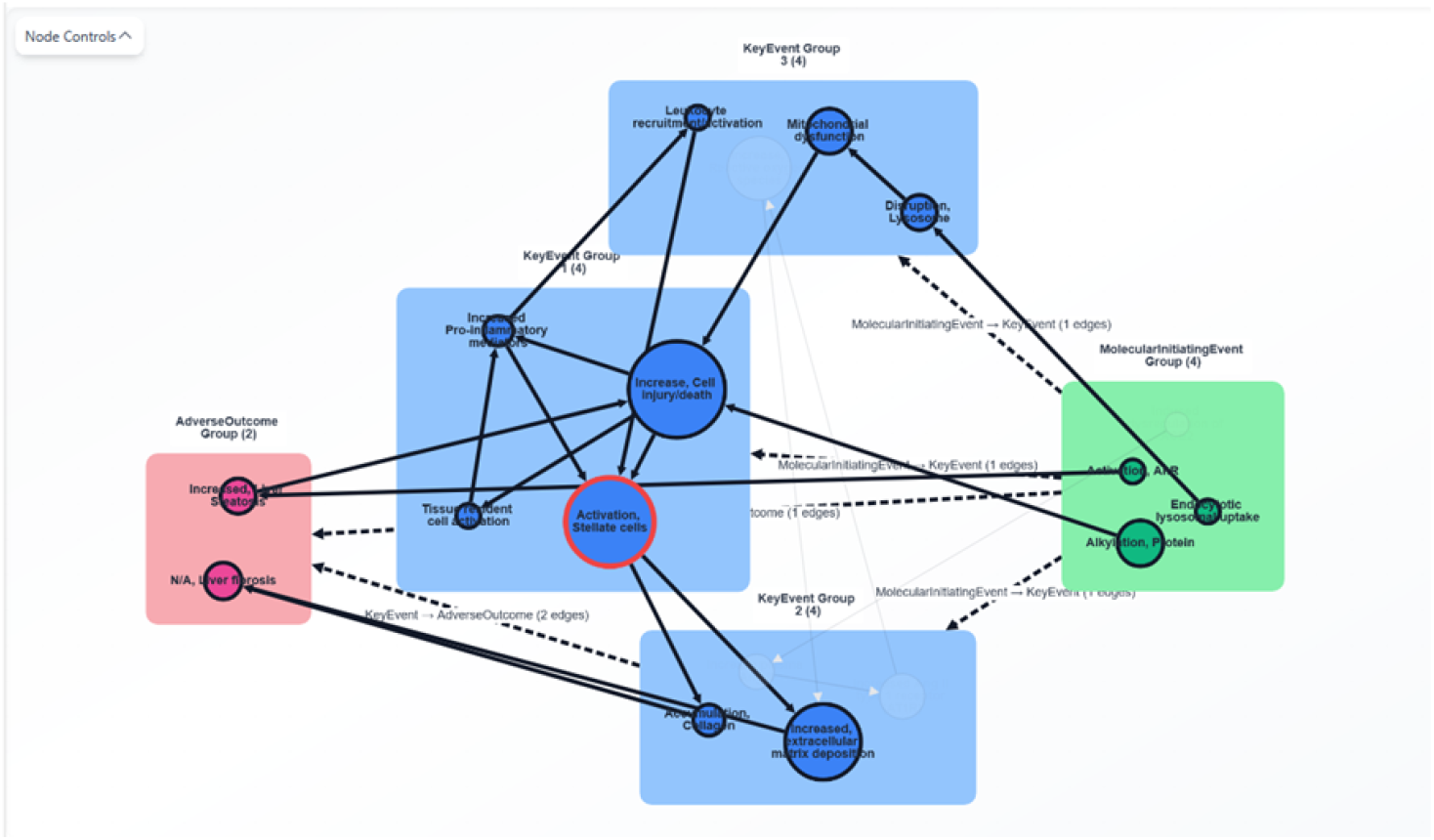
Liver Fibrosis AOP network

### 1.2 State-of-the-Art and Existing Limitations

This need has spurred the development of several specialized AOP visualization and analysis platforms. The AOP-Wiki provides a basic, native network browser, but its static nature and limited interactivity constrain its utility for in-depth analysis.1 More advanced third-party tools have emerged to fill this gap, each representing a step forward in capability.

- AOP-helpFinderJaylet et al. [2025]: This platform integrates text mining of the PubMed database with network visualization using Natural Language Processing (NLP) to identify potential linkages between stressors and KEs from the scientific literature. Its development underscores the importance of augmenting curated AOP knowledge with evidence from broader data sources to support AOP construction.1 The problem of this systems is not full interactive system and it requires initial email and search results are sent as links to personal email and then various metainformation accessible once the network is accessed. However, it cannot design a way to include multiple AOP pathways at once and see overlapping cross-pathway nodes with MIE and AOs.
- AOP-networkFinder Yarar et al. [2025]: This versatile tool focuses on reconstructing and visualizing AOP networks directly from the AOP-Wiki Resource Description Framework (RDF) endpoint. It emphasizes user control, providing features for network filtering, cleaning, and the merging of similar KEs using fuzzy logic, while adhering to FAIR (Findable, Accessible, Interoperable, and Reusable) data principles.
- Wiki Kaptis and Biovista Vizit Biovista [2025]: These platforms focus on enhancing the visualization experience and integrating AOP knowledge into broader biological contexts. Wiki Kaptis, a collaboration between Lhasa Limited Limited [2025]and the AOP-Wiki, excels at displaying the network context of a single KE of interest. 1 Biovista Vizit connects AOP components to extensive knowledge bases of genes, diseases, and pathways, allowing users to explore AOPs within a larger systems biology framework.

The progression of these tools reflects the maturing ambition of the AOP framework. The initial phase focused on documenting individual pathways, which was served by simple viewers. The subsequent phase recognized networks as the functional unit of prediction, leading to tools such as AOP-networkFinder[5] that could reconstruct these networks from underlying data. However, despite these significant advances, the current generation of tools still faces several key limitations that hinder their full potential which are :

- Scalability: Many platforms struggle to render and maintain interactive performance with the increasingly large and dense networks emerging from the AOP-Wiki.
- Standardization: A lack of consistent data formats and analytical capabilities across platforms makes it difficult to compare results or create integrated workflows.
- Representation of Nuance: The representation of temporal dynamics (the timing of events) and the weight of evidence or uncertainty for KERs remains rudimentary, which is a critical gap for regulatory applications.
- Analytical Depth: While existing tools are proficient at visualizing and reconstructing networks, they offer limited capabilities for deep, contextual interrogation of what the network means in a specific biological or chemical context.

### 1.3 Hypothesis and Contribution

This study introduces the AOP Network Viewer, a novel software platform engineered to overcome these limitations and represent the next logical step in the evolution of AOP analysis tools. We hypothesize that by combining a highly scalable and interactive visualization environment with a first-of-its-kind AI-powered contextual analysis engine, we can transform the AOP network from a static object of study into a dynamic and interactive partner in scientific discovery. The AOP Network Viewer is designed not only to display what is known but also to help researchers and regulators ask sophisticated questions and generate new, testable hypotheses directly from the network structure. Its primary contribution is the integration of a Large Language Model (LLM)-based AI Analysis Panel that moves beyond simple reconstruction to enable deep conversational inquiry into the toxicological, pharmacological, and regulatory significance of complex pathway networks. When a user requests an AI analysis, the system must incorporate the node ontology term, identifier, and any secondary metadata into the prompt to generate a more precise and mechanistically accurate interpretation.

## 2 Materials and Methods

This section provides a detailed description of the AOP Network Viewer’s design, architecture, and core functionalities, with sufficient detail to allow for a thorough understanding of its implementation and to support the reproducibility of its conceptual approach.

### 2.1 Network Analysis of AOP Graph

When a cell encounters a toxic substance, it does not simply “break” but embarks on a complex journey through interconnected biological pathways that can lead to disease, dysfunction, or death. Understanding this journey has been one of the greatest challenges in toxicology, but recent advances in network science have finally illuminated the hidden architecture of biological harm. The AOP network structure suggests novel therapeutic strategies. What we understand Rather than addressing individual adverse outcomes, drugs can target the central nodes that control multiple pathways. Antioxidants, mitochondrial protectants, and anti-inflammatory agents have emerged as potential “network drugs” that can provide broad protection against chemical toxicity.

To understand the mechanism, we computed the betweenness centrality scores for each node Kitsak et al. [2007]. The analysis highlighted several nodes with elevated betweenness centrality, including “Oxidative Stress,” “Increase, Reactive oxygen species,” “Mitochondrial dysfunction,” and “Increase, Cell injury/death,” Increase, Cell injury/death. While such centrality metrics are frequently used to identify hub events or potential key mediators in biological networks, this approach has certain limitations in AOP networks. Specifically, these highly central nodes do not consistently provide clear information regarding their upstream or downstream positions within an AOP. Centrality metrics, such as betweenness, reflect the extent to which a node lies on the shortest paths between other nodes but do not explicitly capture the hierarchical directionality or event progression inherent to AOP structures. Consequently, while central nodes may represent common or critical mechanistic events, their identification does not always translate to a mechanistic understanding of the causal sequence or biological context. This complicates efforts to pinpoint where in the pathway critical effects are initiated or amplified, particularly when addressing regulatory or experimental priorities. Incorporating approaches such as pathway tracing, event annotation by biological compartment, or integration with temporal or quantitative data may be required to resolve where these key events occur within the upstream-to-downstream continuum of the AOP, ultimately enhancing its biological interpretability and regulatory utility.

With 1,499 distinct biological events connected by 2,130 relationships, forming what can be thought of as a “biological internet” of toxicological responses. Despite this complexity, the network shows a sparse connectivity pattern (density = 0.0009). The network fragments into 163 connected components, indicating that toxicological responses occur through multiple, semi-independent biological domains—much like different neighborhoods in a vast city, each with its own character and function. Within this network, three distinct classes of biological events: 1,067 Key Events The molecular and cellular processes that propagate toxicological signals, 232 Molecular Initiating Events- The initial chemical-biological interactions that trigger adverse responses and 200 Adverse Outcomes - The final manifestations of toxicity at tissue, organ, and organism levels. This 5:1:1 ratio reveals that for every adverse outcome, there are approximately five intermediate biological steps and one initiating chemical interaction—highlighting the complex amplification cascade that transforms molecular perturbations into organismal harm. Our network analysis identified oxidative stress as the single most critical node in the entire toxicological landscape, with a betweenness centrality of 0.036—making it 1.4 times more central than the second-most important node. This finding provides quantitative evidence for what toxicologists have long suspected: oxidative stress serves as the “master conductor” orchestrating a vast symphony of biological dysfunction. Oxidative stress doesn’t work alone. Our analysis revealed a tight triumvirate of cellular damage mechanisms: Oxidative Stress (centrality: 0.036) - The central coordinator Reactive Oxygen Species(centrality: 0.025) - The molecular messengers Mitochondrial Dysfunction(centrality: 0.024) - The energy crisis together, these three nodes account for nearly 40% of all critical pathway traffic in the network, representing the cellular “perfect storm” that underlies most toxicological responses.

**Figure 2:**
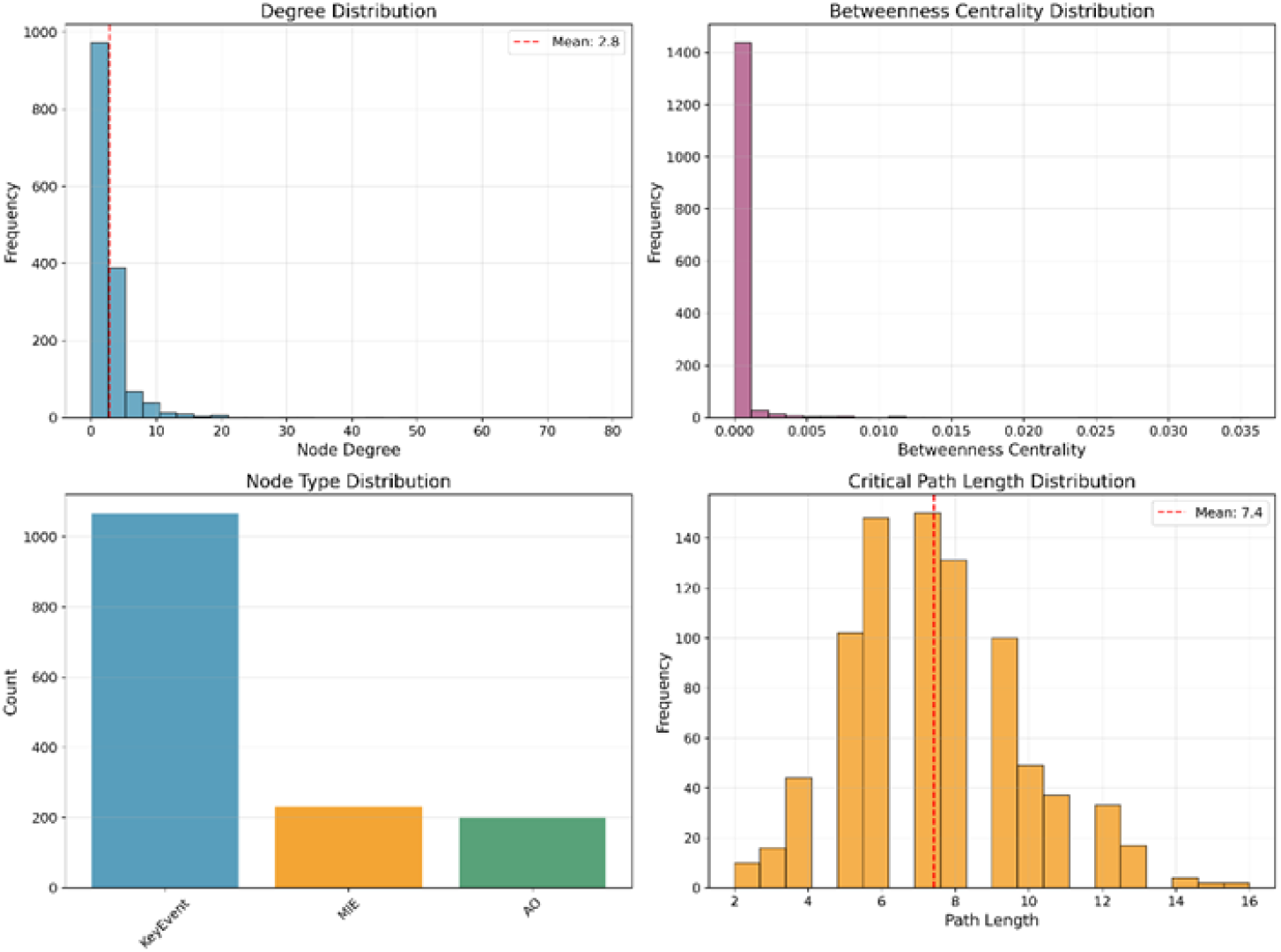
Network Metrics

**Figure 3:**
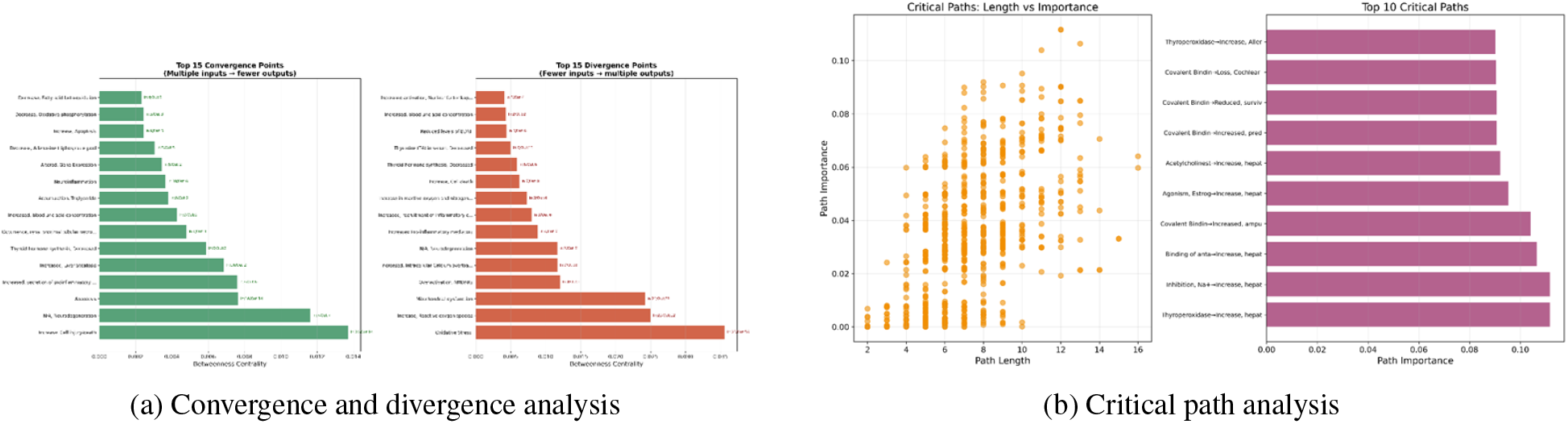
Different Import nodes highlighted

Following oxidative stress in importance, we identified a cascade of increasingly severe cellular responses - Cell Injury/Death(centrality: 0.014) - the point of no return, NMDAR Overactivation (centrality: 0.012) - Neurological crisis, Calcium Overload (centrality: 0.012) - Cellular electrical storm, Neurodegeneration (centrality: 0.012) - Nervous system failure. This hierarchy reveals how chemical exposures progress from subtle molecular perturbations to catastrophic cellular failure, with each step amplifying the damage of the previous one. Our analysis identified 845 critical pathways connecting molecular initiating events to adverse outcomes, with journey lengths averaging 7.4 biological steps. Each critical pathway calculation includes- MIE (Molecular Initiating Event),AO (Adverse Outcome),Complete sequence of biological events, Number of steps in the pathway and Number of steps in the pathway. The five most critical pathways are:

- Thyroid Disruption Highway (Importance: 0.112) Starting with thyroperoxidase inhibition, this 12-step pathway leads inexorably to hepatocellular adenomas and carcinomas. The pathway reveals how endocrine disruption Remarkably, three of the five most critical pathways converge on the same adverse outcome: hepatocellular adenomas and carcinomas. This convergence suggests that the liver serves as a “toxicological sink” the final common pathway for diverse chemical insults, making liver cancer a sentinel endpoint for chemical safety assessment. We identified 360 convergence points biological nodes where multiple toxic pathways merge, potentially creating additive or synergistic effects. The most critical convergence point is cell injury/death, where 20 different pathways converge into 14 downstream consequences. This convergence pattern reveals “toxicological perfect storms” biological situations where multiple chemical exposures could combine to overwhelm cellular defenses. The top convergence points are Cell Injury/Death, Neurodegeneration, Apoptosis, Pro-inflammatory Mediators, Liver Steatosis. Equally important are the 304 divergence points single biological events that trigger multiple downstream consequences. Oxidative stress emerges as the ultimate divergence point, with 23 inputs amplifying into 56 different downstream effects—a 2.4-fold amplification that explains why oxidative stress is so central to toxicological responses. The pattern of divergence reveals biological “amplifier circuits” Oxidative Stress: 23 inputs -> 56 outputs (2.4x amplification), Reactive Oxygen Species: 21 inputs -> 22 outputs (1.0x relay),Mitochondrial Dysfunction: 21 inputs -> 27 outputs (1.3x amplification). These pathways converge on critical cellular fate switches, including p53/p21-mediated cell cycle arrest, NF-KB inflammatory responses, and caspase-mediated apoptosisDi Carlo and Sorrentino [2024]. Chemical designers can use this network map to avoid molecular features that trigger the most critical pathways. The analysis suggests that chemicals should be designed to:
  – Minimize oxidative potential
  – Avoid mitochondrial targeting
  – Prevent non-specific protein binding
  – Limit endocrine disruption potential

### 2.2 Network Visualization and Layout Algorithms

The utility of a network viewer is fundamentally determined by its ability to present complex information in a clear, interpretable manner. The AOP Network Viewer provides multiple layout algorithms, allowing users to select the representation best suited to their analytical question. To maintain a fluid user experience, several performance optimization techniques are employed. Adaptive level-of-detail rendering adjusts the visual complexity of the network based on the user’s zoom level; when zoomed out, nodes may be represented as simple dots, while zooming in reveals detailed labels and annotations. Intelligent caching and predictive loading preload adjacent regions of the network that a user is likely to explore next, minimizing latency during planning and exploration. The entire interface is built on responsive design principles, ensuring full functionality and readability across devices, from large desktop monitors to tablets. The AOP Network Viewer is designed for active exploration, not passive viewing. Its interface is rich with interactive features that empower users to dissect and analyze networks dynamically.

– **Multi-Pathway Selection and Comparison :** The user’s query is normalized and matched against node labels and, optionally, ontology fields using an exact or controlled-fuzzy matching algorithm. By default, ontology-based term expansion is disabled to minimize noise, but it can be enabled by the user. The nodes matched in the initial search seed a set of AOPs. The full node and edge content of these AOPs are then compiled into a candidate graph. A cross pathway property is computed for each node, marking those that belong to more than one AOP. The network is then expanded to include every AOP that contains at least one of these cross-pathway nodes, ensuring that all relevant contexts are captured.
– **Advanced Filtering and Search :** Navigating a network with thousands of nodes requires powerful filtering and search tools. The viewer includes a multi-faceted search interface that supports both exact-match and fuzzy (e.g., Levenshtein distance) searching for KEs, stressors, or genes.1 A hierarchical filtering system allows users to dynamically subset the visible network based on various criteria, such as OECD Nymark et al. [2024] endorsement status, evidence level for KERs, or specific biological ontology terms (e.g., “show only pathways related to inflammation”).
– **Data Integration and Export :** To ensure analytical currency and interoperability, the system features real-time data synchronization with the AOP-Wiki’s backend database, ensuring that any new or updated AOPs are immediately available within the viewer. Recognizing that analysis often continues in other software, the viewer provides comprehensive export capabilities. Users can export the current network view as a high-resolution image (PNG, SVG) suitable for publications. More importantly, they can export the underlying graph data and associated attributes in structured formats like JSON or CSV. This includes a dedicated export option for Cytoscape Burgoon [2017] Shannon et al. [2003], a widely used platform for biological network analysis Pavlopoulos et al. [2011], allowing for seamless integration into existing bioinformatics workflows and promoting FAIR data principles.

**Figure 4:**
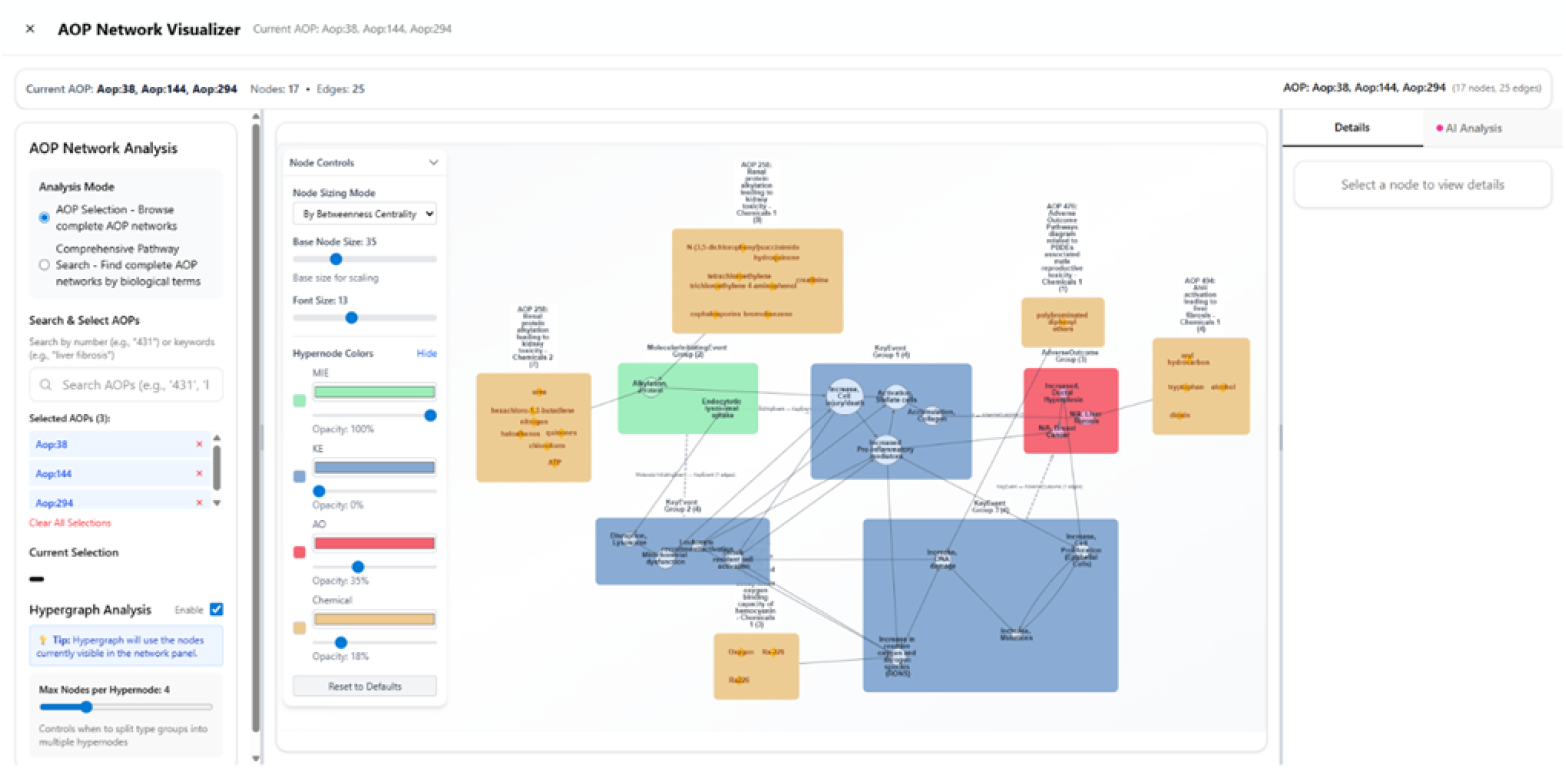
AOP Net UI

**Figure 5:**
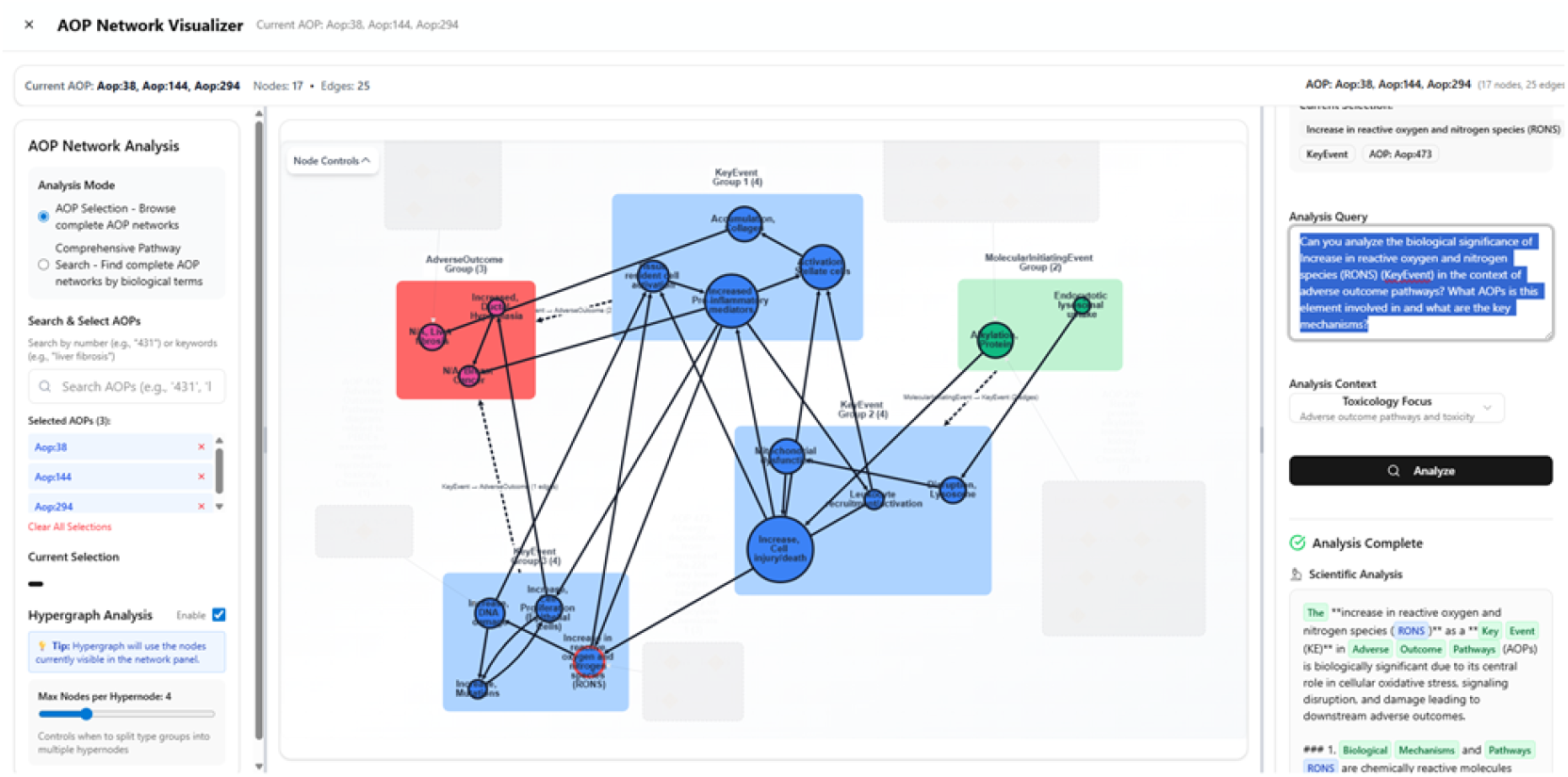
AOP Net with LLM integration
– **The AI Analysis Panel:** LLM-Powered Contextual Inquiry : The most significant innovation of the AOP Network Viewer is the AI Analysis Panel, a feature that integrates a powerful Large Language Model (LLM) directly into the analysis workflow. This transforms the tool from a data visualizer into an interactive research assistant, capable of synthesizing information, answering complex questions, and generating hypotheses. This approach addresses a major challenge in systems biology: bridging the gap between the structured, graphical representation of a pathway and the vast, unstructured knowledge contained within the scientific literature[8,9,10]. The system’s design directly confronts the known limitations of general-purpose LLMs, such as lack of domain-specific context and the potential for “hallucination,” through a robust and carefully engineered prompt architecture[27].

The prompt design philosophy ensures context-aware responses by dynamically building queries that include the user’s selection state (nodes, edges, pathways), current AOP context, and analysis type preferences. Within the AI’s generated response, key biological entities such as genes, diseases, chemical compounds, and pathways are automatically recognized and highlighted with distinct, color-coded badges. This allows for rapid visual parsing of the information. Each of the 7 modes (General, Pathway, Mechanism, Toxicology, Disease, Regulatory, Drug Analysis) applies domain-specific prompt enhancements, with the Drug Analysis mode being the most sophisticated—automatically expanding queries to include therapeutic targets, regulatory status, and personalized medicine approaches, while the “More” analysis provides a second-tier prompt focusing on clinical translation and real-world application of the pathway knowledge.

**LLM Context**

Can you analyze the biological significance of Increase in reactive oxygen and nitrogen species (RONS) (KeyEvent) in the context of adverse outcome pathways? What AOPs is this element involved in and what are the key mechanisms?

**LLM Reasoning**

The increase in reactive oxygen and nitrogen species (RONS) as a Key Event (KE) in Adverse Outcome Pathways (AOPs) is biologically significant due to its central role in cellular oxidative stress, signaling disruption, and damage leading to downstream adverse outcomes. Mitochondria are a critical source and target of RONS; regulatory proteins such as PHLPP1 influence region-specific mitochondrial dynamics linked to oxidative stress responses.

- Biological Mechanisms and Pathways: RONS are chemically reactive molecules derived from oxygen and nitrogen, including free radicals like superoxide (O2•), hydroxyl radicals (•OH), and non-radical species like hydrogen peroxide (H2O2) and nitric oxide (NO). They are generated endogenously through mitochondrial electron transport, enzymatic activity (e.g., NADPH oxidases, nitric oxide synthases), or exogenously through xenobiotic metabolism.
  * Oxidative Stress Induction: Excess RONS overwhelm cellular antioxidant defenses, resulting in oxidative stress, which causes damage to lipids, proteins, and DNA.
  * Mitochondrial Dysfunction: RONS can trigger mitochondrial permeability transition and fission, leading to energy metabolism disruption and apoptosis pathways activation.
  * Inflammation Activation:RONS activate redox-sensitive transcription factors (e.g., NF- KB), inducing pro-inflammatory gene expression.
  * Cell Signaling Perturbation: RONS modulate kinase and phosphatase activities, affecting signaling cascades regulating cell proliferation, differentiation, and death[4].
- Molecular Interactions and Processes: At the molecular level, increased RONS lead to:
  * Lipid Peroxidation: Formation of reactive aldehydes that further propagate cellular damage.
  * Protein Oxidation and Nitrosylation: Affecting enzyme activity and structural integrity.
  * DNA Damage:Induction of base modifications, strand breaks, mutations contributing to genomic instability.
  * Cross-talk with Immune Cells: RONS released by neutrophils and other immune cells exacerbate tissue damage and thrombosis risk in inflammatory diseases[3].
- Toxicological Significance: The perturbation of redox balance via increased RONS is a pivotal early event in many toxicological pathways:
  * Cytotoxicity and Cell Death: RONS accumulation can initiate apoptosis, necrosis, or ferroptosis (an iron-dependent form of cell death characterized by lipid peroxidation)[2].
  * Organ Toxicity:Elevated RONS are implicated in acute kidney injury, neurodegeneration, cancer progression, and cardiovascular diseases due to oxidative damage and inflammation[4][5].
  * Disease Initiation and Progression:Chronic oxidative stress can result in mutations and dysregulated cell signaling, contributing to carcinogenesis and chronic inflammation-related pathologies.
- Involvement in AOPs and Literature Evidence While the query includes the AOP identifier **AOP:473**, comprehensive details on its contents were not retrieved. However, increase in RONS is known to be a **common key event in numerous AOPs**, where it acts as a central mediator linking molecular-level stressors to adverse outcomes. Examples of ROS/RONS involvement in AOPs include:
  * AOPs leading to hepatotoxicity via mitochondrial dysfunction.
  * Neurotoxicity pathways involving astroglial mitochondrial fission regulation by PHLPP1 linked to ROS generation[1].
  * Ferroptosis-related AOPs where RONS mediate lipid peroxidation leading to cell death[2].
  * Inflammatory cardiovascular AOPs where RONS promote endothelial injury and thrombosis through immune cell activation[3].

Key recent literature highlights the fine regulation of RONS in health and disease signaling pathways, such as nuclear receptors modifying RONS generation to influence acute kidney injury and other diseases[4], emphasizing their dual role as physiological signaling molecules and pathological mediators.

– **Drug Analysis Mode:** This is the most sophisticated mode, designed for pharmaceutical and therapeutic research. It automatically expands a user’s query by appending focused prompts about “drugs modulating the selected MIE/KE pathways, FDA-approved compounds targeting these entities, known drug-target interactions, and potential drug repurposing opportunities.” This transforms a simple selection into a comprehensive pharmacological investigation.
– **Toxicology Focus Mode:** This mode is tailored for risk assessment. It enhances the query to specifically request information on “the biological mechanisms, key molecular interactions, overall toxicological significance, and known regulatory relevance of the selected pathways.”
– **Pathway Analysis Mode**: Focuses on biological pathway flow and causal relationships within AOPs. Analyzes upstream molecular initiating events (MIEs), intermediate key events (KEs), and downstream adverse outcomes (AOs) with emphasis on adjacency relationships and confidence levels. Identifies critical pathway nodes, branching points, and regulatory bottlenecks that determine biological response patterns.
– **Mechanism Study Mode :** Provides detailed molecular-level explanations of biochemical processes and protein interactions. Examines specific molecular targets, enzyme activities, receptor binding, and cellular signaling cascades that drive each key event. Integrates ontology terms (GO, MeSH) to explain precise molecular mechanisms underlying toxicological responses.
– **Disease Context Mode :** Connects AOP pathways to human pathology and clinical outcomes by linking adverse outcomes to specific diseases, syndromes, and health endpoints. Translates molecular events into clinically relevant phenotypes, helping researchers understand how environmental exposures lead to observable health effects and disease manifestations in human populations.

Every query to the AI panel is composed with a three-layer architecture that ensures answers are precise, relevant, and scientifically rigorous. A permanent system instruction anchors the model as an expert in toxicology, systems biology, pharmacology, and regulatory science, directing responses toward evidence-based, domain-specific reasoning. A dynamic context injection step then embeds the user’s current graphical selections such as specific KEs, KERs, and AOP titles into the prompt, ensuring the model’s analysis targets the user’s exact focus. Finally, enhanced query processing wraps the user’s natural language question in a context-mode template that prescribes the type of analysis required, producing structured, actionable outputs even from imprecise queries. Combined, these layers convert simple prompting into programmatic prompt engineering, tightly constraining the LLM’s broad knowledge to the task at hand.

## 3 Results

### 3.1 Case Study 1: Elucidating Kidney Failure Networks for Alternative Testing Strategy Development

The assessment of nephrotoxicity (kidney toxicity) is a critical challenge in drug development and chemical safety assessment. The kidneys are highly susceptible to damage from a wide range of chemicals due to their high bloodflow, concentrated filtering processes, and active transport mechanisms. Traditional animal testing for nephrotoxicity is slow, expensive, and raises ethical concerns. The AOP framework provides a powerful alternative by elucidating the mechanistic pathways leading to kidney failure, enabling the development of more targeted and efficient in vitro testing strategies. In this case study, the AOP Network Viewer was used to analyze a complex network of AOPs associated with kidney failure. The goal was to identify critical “hub” key events (KEs) that could serve as focal points for a tiered in vitro testing strategy.

Using the AOP Network Viewer, differentent MIEs implicated in nephrotoxicity were identified, for example: inhibition of mitochondrial DNA polymerase gamma (impairs mitochondrial replication),receptor-mediated endocytosis with lysosomal overload (drug accumulation lysosomes), and covalent protein binding/alkylation (cell dysfunction and immune activation). Despite disparate initiators, the pathways converged on a small set of hub key events notably mitochondrial dysfunction, oxidative stress, and tubular necrosis which together constitute the mechanistic backbone leading to kidney failure. Guided by these hubs, we propose a tiered testing strategy: Tier 1 : high-throughput in vitro screens for mitochondrial function and oxidative stress to rapidly flag chemicals of concern; Tier 2 : targeted assays to probe specific upstream MIEs and confirm mechanism; and Tier 3 : focused in vivo or advanced organoid studies for high-priority compounds. This network-informed workflow prioritizes biologically central assays, reduces reliance on broad in vivo testing, and accelerates mechanistic, regulatory relevant safety assessment.

**Figure 6:**
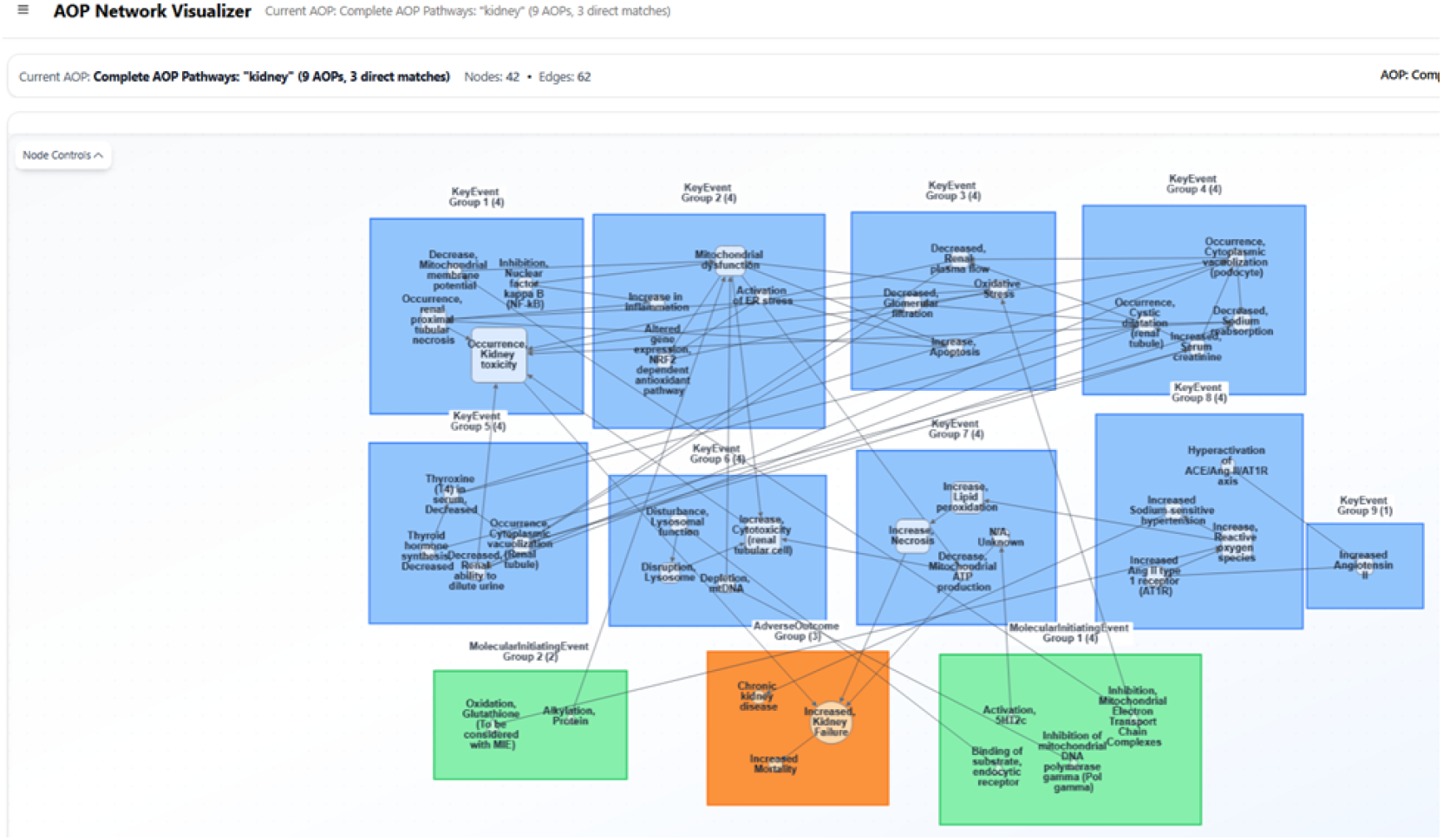
(AOP for kidney toxicity)

**Figure 7:**
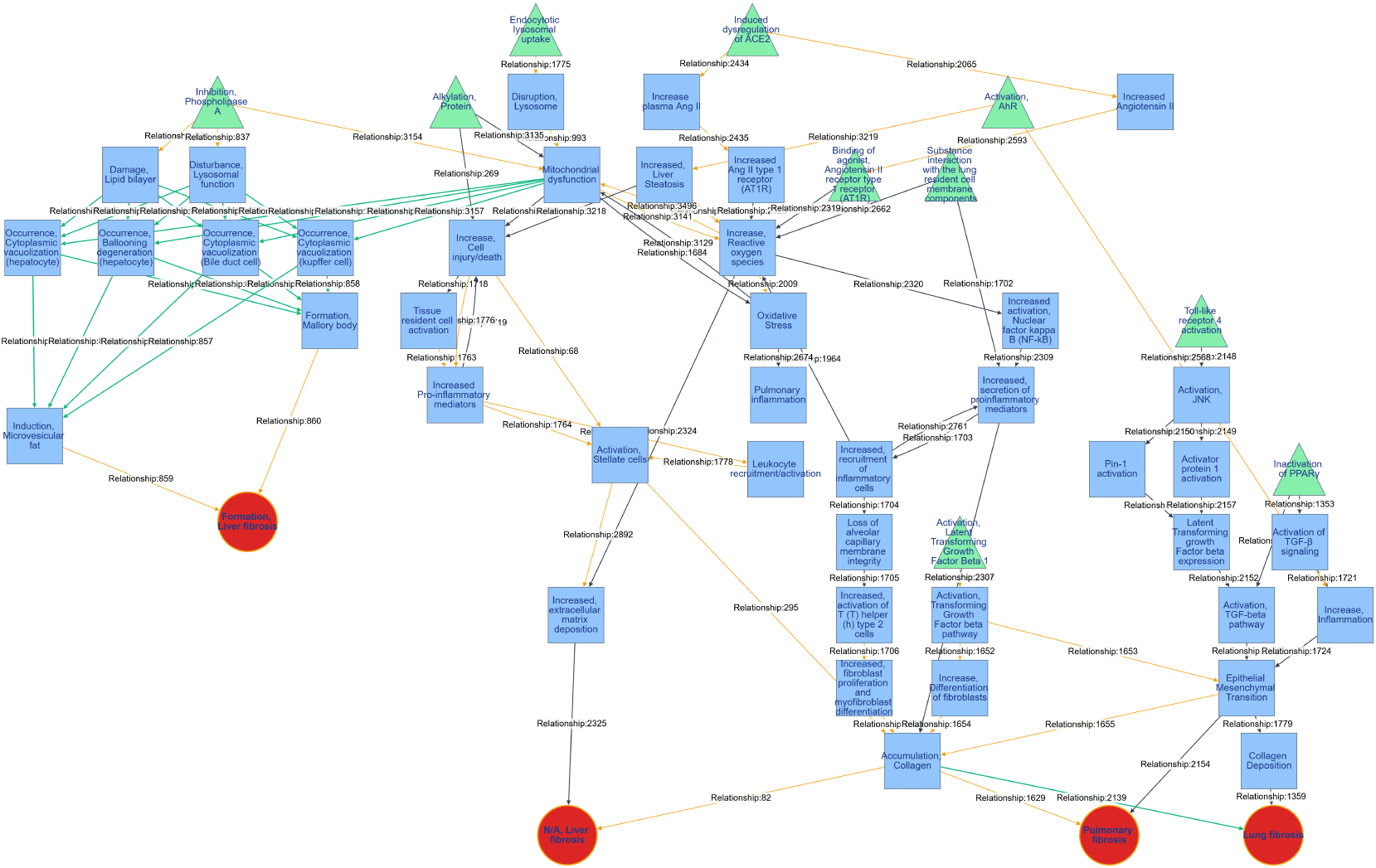
AOP for Liver Fibrosis and Lung Fibrosis with Key event Overlap of Accumulation of collagen

## 4 Discussion

The development of the AOP Network Viewer represents a significant step forward in the field of computational toxicology. This work’s principal achievement is the creation of a next-generation software platform that not only addresses the key technical limitations of previous tools but also introduces a new paradigm for interacting with and analyzing complex biological knowledge. The integration of a scalable, interactive visualization environment with a powerful, context-aware AI engine transforms the process of AOP analysis from one of passive data consumption to one of active, conversational scientific discovery. The primary novelty lies in the AI Analysis Panel, which empowers users to move beyond observing network structures to deeply interrogating their biological, toxicological, and therapeutic meaning.

The AI Analysis Panel in the AOP Network Viewer represents a more sophisticated and targeted application of this technology. Its three-layer prompt architecture is a deliberate engineering solution designed to mitigate the known weaknesses of LLMs, such as their tendency to “hallucinate” or provide generic, out-of-context answers when applied to specialized domains. By programmatically injecting specific, user-selected network context (a form of real-time RAG) and framing queries with domain-specific instructions, the system constrains the LLM to function as a reliable scientific assistant. This approach aligns with state-of-the-art research on enhancing LLM reasoning for biological pathways, such as the proposed Path Finder search, which similarly leverages subgraph navigation within a larger knowledge graph to improve reasoning about the effects of perturbations. Our implementation provides a practical, user-friendly interface to this powerful concept, making advanced AI-driven analysis accessible to the broader toxicology community.

### 4.1 Addressing the Limitations of Previous Tools

The design of the AOP Network Viewer was explicitly guided by the need to overcome the documented limitations of existing platforms.

- **Scalability and Performance:** The client server architecture and graph database backend directly address the scalability issues that cause performance degradation in other tools when handling large networks.
- **Interactivity and Analytical Depth:** While tools like AOP-networkFinder Yarar et al. [2025]offer excellent reconstruction capabilities, and Biovista Vizit provides broad knowledge integration, the AOP Network Viewer combines these strengths with a higher level of interactivity, including dynamic multi-pathway analysis.
- Contextual Analysis: The AI Analysis Panel provides a level of deep, contextual inquiry that is absent in other tools. While AOP-helpFinder uses NLP for literature mining to build AOPs, our system uses an LLM to interpret existing AOP networks in the context of pharmacology, disease, and regulation, representing a significant leap in analytical capability.
- **Representation of Nuance:** Although still a challenge, the AI-driven approach allows for a more nuanced discussion of evidence and uncertainty. The LLM can summarize and present the weight of evidence described in the literature, moving beyond the simple graphical links of other viewers.

### 4.2 Generating Hypothesis for new modalities

From Figure 6 tools enabling graph and networks can be used to design new modalities like for example the biological significance of AT1R agonist binding in liver fibrosis adverse outcome pathways represents a compelling case study for how computational tools and systematic evidence evaluation can identify gaps in current AOP frameworks and generate novel mechanistic hypotheses. While extensive evidence demonstrates that angiotensin II binding to AT1R plays a central role in liver fibrosis progression through well-characterized pathways including JAK2/RhoA activation, TGF-*β* upregulation, and hepatic stellate cell activation, no formal AOP currently exists with AT1R agonist binding as the molecular initiating event (MIE) for liver fibrosis. This represents a significant knowledge gap, as the existing liver fibrosis AOPs focus on protein alkylation (AOP 38) or endocytic lysosomal uptake (AOP 144) as primary MIEs, while AT1R-mediated mechanisms appear only as key events in related pathways like ACE2 inhibition leading to liver fibrosis (AOP 383). The evidence base strongly supports AT1R binding as a legitimate MIE candidate - it represents the initial molecular interaction between angiotensin II and its cognate GPCR, triggers multiple downstream pro-fibrotic cascades through JAK2 phosphorylation and RhoA activation, and demonstrates clear dose-response relationships with therapeutic intervention studies showing AT1R antagonists effectively prevent liver fibrosis Jaeger et al. [2017].

### 4.3 Limitations of the Current Work

Despite its advancements, it is crucial to acknowledge the limitations of the AOP Network Viewer to guide its appropriate use and future development.

- **Data Dependency:** The adage “garbage in, garbage out” applies directly. The quality of both the visual network representation and the AI-generated analysis is fundamentally dependent on the accuracy, completeness, and curation quality of the underlying data in the AOP-Wiki and the literature the LLM was trained on. Inaccuracies or gaps in the AOP-Wiki will be reflected and potentially amplified by the tool. There are several AOPs which are not correctly formatted and llm tools can help to make it correct and easier to interpret for example AOP:423 regarding Activation of PARP1. Instead of a logical cascade, it is a jumbled collection of terms. The correct pathway begins with the Molecular Initiating Event (MIE 1115), an increase in reactive oxygen species (ROS). This oxidative stress then leads to the first Key Event (KE 1194), an increase in DNA damage. This damage, in turn, serves as the signal for the activation of PARP1 (MIE 2030). The hyperactivation of PARP1 is a crucial step that leads directly to mitochondrial dysfunction (KE 177), primarily by depleting the cell’s energy and creating PAR polymers. A specific and critical consequence of this dysfunction is the releasing of Apoptosis-Inducing Factor (AIF) (KE 2031) from the mitochondria. This highlights the main sequencing error in the original list: the events were not linked by cause and effect. The confusion happens because “heart failure” (https://aopwiki.org/events/1535), “Increased, Kidney Failure”, “Pulmonary fibrosis”,can be viewed in two ways: it’s a complex process of organ-level dysfunction, but it’s also a clinically defined disease that is adverse in itself. An AOP must be a linear path, so an event can’t be both a step along the way (KE) and the final destination (AO).
- **AI Oversight and Verification:** The AI Analysis Panel is a powerful hypothesis-generation engine, not an arbiter of truth. All AI-generated insights, especially novel hypotheses about drug repurposing or mixture toxicity, must be treated as preliminary and require rigorous validation by human experts and subsequent experimental testing. The risk of LLM bias or subtle factual errors (hallucinations), while mitigated by our prompt architecture, can never be fully eliminated.30 The clickable reference links are a key feature designed to empower users in this critical verification step. A single, dense diagram Figure 5, even if mechanistically sound, can be overwhelming. A more effective approach is to deconstruct the pathways into their core components and present the causal relationships in a structured format, such as a table. This approach forces a detailed, narrative explanation of each KER, providing the depth and clarity that a diagram alone cannot. For example AOP 382 the link between AT1R binding and increased reactive oxygen species (ROS) is missing the key enzyme NADPH oxidase, which is the primary source of ROS in this pathway. Similarly, the connection between ROS and NF-kB activation lacks the crucial intermediate step involving the inactivation of I-kB, which normally holds NF-kB in check. Lastly, the AOP could be strengthened by listing the specific pro-inflammatory mediators, such as TNF-*α* and IL-6, that are secreted as a result of NF-kB activation, providing a more tangible link to inflammation. To improve the AOP, these missing mechanistic details should be added as new key events or as more descriptive text within the existing ones, transforming it from a simple flowchart into a robust, evidence-based biological pathway.
- **Static Knowledge Cutoff:** The core knowledge of the LLM is limited to the data it was trained on, which has a specific cutoff date. The context injection mechanism provides real-time information about the network structure itself, but the LLM’s ability to interpret that structure is based on its past training. This means it may not be aware of the very latest research findings not yet incorporated into its model, a common challenge for all LLM applications in fast-moving scientific fields.
- **Limitations of AOP Networks:** The accuracy of critical path identification is fundamentally constrained by the scope and completeness of existing toxicological data. Since AOP networks rely on available mechanistic knowledge, gaps in the scientific literature directly translate to potential blind spots in critical path determination. This limitation is particularly significant given that the AOP knowledgebase remains incomplete, with many pathways still undiscovered or inadequately annotated. Current critical path identification approaches remain relatively loosely defined, with different stakeholders interpreting “critical” through various lenses - whether from investigational, biological, or regulatory perspectives. The development of quantitative AOPs (qAOPs) Conolly et al. [2017] may be necessary to establish more stringent and objective criteria for critical path determination. Critical paths may not necessarily follow single AOPs but rather emerge through the assembly and consideration of interactions between multiple AOPs. This complexity requires sophisticated analytical approaches that can effectively integrate across interconnected pathway networks while maintaining biological relevance. A major challenge lies in determining whether critical paths will be conserved across different species. Even when AOPs are deemed relevant to specific taxonomic domains, the relative importance of different pathways may vary significantly between species. This creates substantial uncertainty for cross-species extrapolation in risk assessment, as a pathway that appears critical in one test species may not hold the same significance in the target species of concern. sectionConclusions and Future Directions The development and application of advanced AOP network viewers represent a transformative advancement for computational toxicology. Our comprehensive analysis reveals that toxicological responses operate through a “biological internet” a sparse but highly organized network where oxidative stress serves as the central hub, coordinating responses across multiple biological systems. The network’s architecture creates both vulnerability (through amplification cascades) and opportunity (through identifiable intervention points). This network perspective transforms our understanding of chemical toxicity from a collection of isolated adverse effects to an integrated system of biological responses. This paper has presented the AOP Network Viewer, a software platform that addresses the critical limitations of previous tools and establishes a new benchmark for the analysis of complex toxicological knowledge. Through a comprehensive analysis of its technical innovations and practical applications, we have demonstrated that this sophisticated network visualization and analysis tool is not merely a useful addition to the toxicological toolkit, but essential infrastructure for the continued evolution of 21st-century toxicology. The key conclusions of this work are fourfold:
  - Technological Maturity: The AOP Network Viewer represents a mature technology ready for widespread adoption. Its scalable architecture, responsive user interface, and diverse analytical features successfully address the major performance and usability limitations of earlier approaches.
  - Transformative AI Integration: The integration of a context-aware LLM via the AI Analysis Panel is a transformative enhancement. It moves beyond simple visualization to enable intelligent interpretation, automated analysis, and predictive hypothesis generation, making deep mechanistic inquiry accessible to a broad range of users.
  - Demonstrated Practical Value: The case studies provide compelling evidence of the platform’s value in critical applications, including alternative testing strategy development, chemical prioritization, and mixture toxicity assessment. These applications show that a network-based approach enhances both the efficiency and the scientific rigor of toxicological assessments.
  - Foundation for Future Innovation: The platform’s modular and extensible design provides a robust foundation for future development, particularly in the promising and critical area of personalized risk assessment.

## 5 Extensibility to Personalized Toxicology

The most significant future direction is the extension of the platform to personalized toxicology and risk assessment. The vision is to move beyond population-average AOPs to individualized pathway models that account for the genetic, environmental, and lifestyle factors known to modulate human susceptibility to toxicants.1 The system’s modular architecture is designed to support this by allowing the dynamic modification of pathway parameters based on individual-level data. For example:

- Genetic Variability: We plan to integrate data on common genetic polymorphisms, such as those in cytochrome P450 enzymes (e.g., CYP2D6), which are critical for chemical metabolism.1 A user could input an individual’s genotype, and the network model would adjust the rates of specific metabolic KEs, providing a personalized prediction of internal dose and downstream pathway activation.
- Environmental and Lifestyle Factors: The model will be extended to incorporate data on co-exposures, nutritional status (e.g., selenium levels affecting oxidative stress pathways), and disease states (e.g., liver disease impairing metabolism), allowing for a more holistic and context-specific risk assessment.

A final, ambitious goal is to enable the AOP Network Viewer to contribute back to the scientific commons. A future version could include functionality that allows users, particularly expert curators, to flag potential new KEs or KERs identified through AI-driven analysis. The system could format these suggestions for semi-automated submission back to the AOP-Wiki, thus creating a virtuous cycle where the tool not only consumes but also helps to generate and refine the community’s collective knowledge base, addressing a key challenge in third-party tool integration. In summary, the AOP Network Viewer system developed in this paper represents a significant step forward in our ability to understand, analyze, and apply mechanistic toxicological knowledge. As it continues to evolve, it promises to play an increasingly central role in advancing the goals of 21st-century toxicology: reducing animal testing, improving the accuracy of safety assessments, and supporting evidence-based regulatory decisions that protect human and environmental health.

## 6 Acknowledgements

## Author Contributions

Conceptualization, methodology AS

## Institutional Review Board Statement

Not applicable. This study did not involve humans or animals.

## Informed Consent Statement

Not applicable. This study did not involve humans.

## Data Availability Statement

The AOP Network Viewer software is a proprietary project. Data used for case studies were derived from publicly available sources, including the AOP-Wiki (aopwiki.org), the EPA’s CompTox Chemicals Dashboard (comptox.epa.gov/dashboard) and https://github.com/abhik1368/aop-visualizer-clean.

## Acknowledgments

The authors wish to thank the developers and curators of the AOP-Wiki and the various third-party tool developers whose work provided the foundation and context for this project. I also want to thank Claud Code to polishing and helping in vibe coding of the frontend of UI and helping in designing the sophisticated features.

